# Genetic basis of body color and spotting pattern in redheaded pine sawfly larvae (*Neodiprion lecontei*)

**DOI:** 10.1101/183996

**Authors:** Catherine R. Linnen, Claire T. O’Quin, Taylor Shackleford, Connor R. Sears, Carita Lindstedt

**Affiliations:** Department of Biology, University of Kentucky, Lexington, KY, 40506; Current affiliation: Department of Biological Sciences, University of Cincinnati, Cincinnati, OH, 45221; Centre of Excellence in Biological Interactions, Department of Biological and Environmental Sciences, University of Jyväskylä, Jyväskylä, Finland, FI-40014

**Keywords:** pigmentation, carotenoids, melanin, genetic architecture, convergent evolution, evolutionary genetics

## Abstract

Pigmentation has emerged as a premier model for understanding the genetic basis of phenotypic evolution, and a growing catalog of color loci is starting to reveal biases in the mutations, genes, and genetic architectures underlying color variation in the wild. However, existing studies have sampled a limited subset of taxa, color traits, and developmental stages. To expand our sample of color loci, we performed quantitative trait locus (QTL) mapping analyses on two types of larval pigmentation traits that vary among populations of the redheaded pine sawfly (*Neodiprion lecontei*): carotenoid-based yellow body color and melanin-based spotting pattern. For both traits, our QTL models explained a substantial proportion of phenotypic variation and suggested a genetic architecture that is neither monogenic nor highly polygenic. Additionally, we used our linkage map to anchor the current *N. lecontei* genome assembly. With these data, we identified promising candidate genes underlying: (1) a loss of yellow pigmentation in Mid-Atlantic/northeastern populations (*Cameo2* and *apoLTP-II/I*), and (2) a pronounced reduction in black spotting in Great-Lakes populations (*yellow, TH, Dat*). Several of these genes also contribute to color variation in other wild and domesticated taxa. Overall, our findings are consistent with the hypothesis that predictable genes of large-effect contribute to color evolution in nature.

## INTRODUCTION

Over the last century, color phenotypes have played a central role in our efforts to understand how evolutionary processes shape phenotypic variation in natural populations (Gerould 1921; Sumner 1926; Fisher and Ford 1947; Haldane 1948; Cain and Sheppard 1954; Kettlewell 1955). More recently, as technological advances have enabled us to link genotypes and phenotypes in non-model organisms (Davey *et al*. 2011; Gaj *et al*. 2013; Goodwin *et al*. 2016; Huang *et al*. 2016), we’ve begun to decipher the genetic and developmental mechanisms underlying naturally occurring color variation as well (True 2003; Protas and Patel 2008; Wittkopp and Beldade 2009; Manceau *et al*. 2010; Nadeau and Jiggins 2010; Kronforst *et al*. 2012). With a growing catalog of color loci, we are starting to assess how ecology, evolution, and development interact to bias the genetic architectures, genes, and mutations underlying the remarkable diversity of color phenotypes in nature (Hoekstra and Coyne 2007; Stern and Orgogozo 2008, 2009; Kopp 2009; Manceau *et al*. 2010; Streisfeld and Rausher 2011; Martin and Orgogozo 2013; Dittmar *et al*. 2016; Massey and Wittkopp 2016; Martin and Courtier-Orgogozo 2017). To make robust inferences about color evolution, however, we require genetic data from diverse traits, taxa, developmental stages, and evolutionary timescales.

Two long-term goals of research on the genetic underpinnings of pigment variation are: (1) to evaluate the importance of large-effect loci to phenotypic evolution (Orr and Coyne 1992; Mackay *et al*. 2009; Rockman 2012; Remington 2015; Dittmar *et al*. 2016), and (2) to determine the extent to which the independent evolution of similar phenotypes (phenotypic convergence) is attributable to the same genes and mutations (genetic convergence) (Arendt and Reznick 2008; Gompel and Prud’homme 2009; Christin *et al*. 2010; Manceau *et al*. 2010; Elmer and Meyer 2011; Conte *et al*. 2012; Rosenblum *et al*. 2014). Addressing these questions will require a large, unbiased sample of pigmentation loci. At present, most of what we know about the genetic basis of color variation in animals comes from a handful of taxa and from melanin-based color variation in adult life stages (True 2003; Protas and Patel 2008; Wittkopp and Beldade 2009; Nadeau and Jiggins 2010; Kronforst *et al*. 2012; Sugumaran and Barek 2016). Another important source of bias in our existing sample of pigmentation loci stems from a tendency to focus on candidate genes and discrete pigmentation phenotypes (Kopp 2009; Manceau *et al*. 2010; Rockman 2012; but see O’Quin *et al*. 2013; Albertson *et al*. 2014; Signor *et al*. 2016; Yassin *et al*. 2016). Here, we describe an unbiased, genome-wide analysis of multiple, continuously varying color traits in larvae from the order Hymenoptera, a diverse group of insects that is absent from our current catalog of color loci (Martin and Orgogozo 2013).

More specifically, our study focuses on pine sawflies in the genus *Neodiprion*. Several factors make *Neodiprion* an especially promising and tractable system for investigating the genetic basis of color variation. First, there is extensive variation in many different types of color traits, and these vary both within and between *Neodiprion* species (Figures 1-2). Second, previous phylogenetic and demographic studies of this genus (Linnen and Farrell 2007, 2008a; b; Bagley *et al*. 2017) enable us to infer directions of trait change and identify instances of phenotypic convergence. Third, because many different species can be reared and crossed in the lab (Knerer and Atwood 1972, 1973; Kraemer and Coppel 1983; Bendall *et al*. 2017), unbiased genetic mapping approaches are feasible in *Neodiprion*. Fourth, a growing list of genomic resources for *Neodiprion*—including an annotated genome and a methylome for *N. lecontei* (Vertacnik *et al*. 2016; Glastad *et al*. 2017)— facilitate identification of causal genes and mutations. And finally, an extensive natural history literature (Coppel and Benjamin 1965; Knerer and Atwood 1973) provides insights into the ecological functions of color variation in pine sawflies, which we review briefly for context.

**Figure 1.**
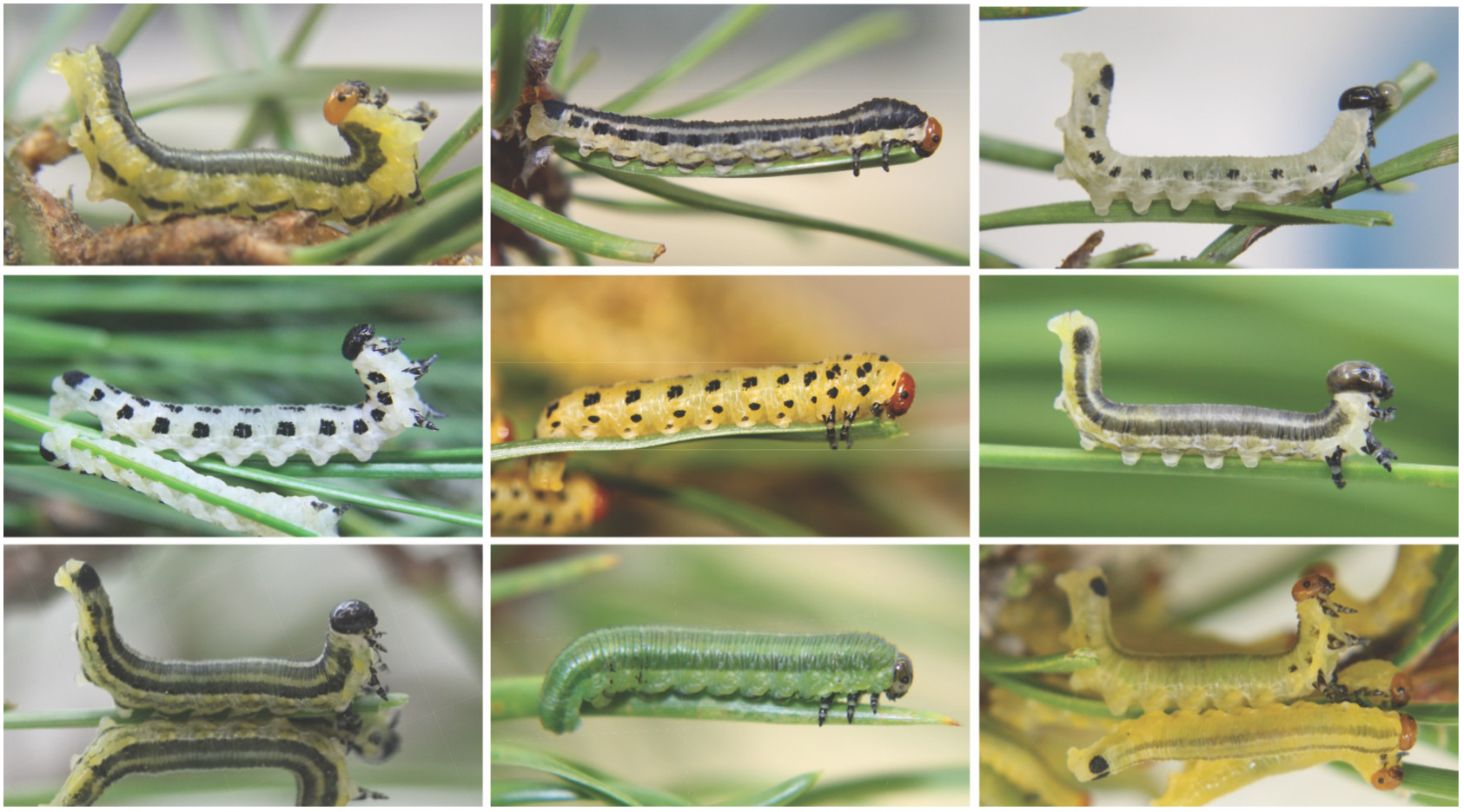
Interspecific variation in *Neodiprion* larval color. Top row (left to right): *Neodiprion nigroscutum, N. rugifrons, N. virginianus*. Middle row (left to right): *N. pinetum, N. lecontei, N. merkeli*. Bottom row (left to right): *N. pratti, N. compar, N. swainei*. Larvae in the first and last columns are exhibiting a defensive “U-bend” posture (a resinous regurgitant is visible in *N. virginianus*, top right). *N. pratti* photo is by K. Vertacnik, all others are by R. Bagley.

Under natural conditions, pine sawfly larvae are attacked by a diverse assemblage of arthropod and vertebrate predators, by a large community of parasitoid wasps and flies, and by fungal, bacterial, and viral pathogens (Coppel and Benjamin 1965; Wilson *et al*. 1992; Codella and Raffa 1993). When threatened, larvae adopt a characteristic “U-bend” posture and regurgitate a resinous defensive fluid (Figure 1), which is an effective repellant against many different predators and parasitoids (Eisner *et al*. 1974; Codella and Raffa 1995; Lindstedt *et al*. 2006, 2011). Although most *Neodiprion* species are chemically defended, larvae vary from a green striped morph that is cryptic against a background of pine foliage to highly conspicuous aposematic morphs with dark spots or stripes overlaid on a bright yellow or white background (Figure 1). Thus, larval color is likely to confer protection against predators either via preventing detection (crypsis) or advertising unpalatability (aposematism) (Ruxton *et al*. 2004; Lindstedt *et al*. 2011).

Beyond selection for crypsis or aposematism, there are a myriad of abiotic and biotic selection pressures that could act on *Neodiprion* larval color. For example, coloration plays diverse ecological roles in insects, including thermoregulation, protection against UV damage, desiccation tolerance, and resistance to abrasion (True 2003; Lindstedt *et al*. 2009; Wittkopp and Beldade 2009). Color alleles may also have pleiotropic effects on other traits, such as behavior, immune function, diapause/photoperiodism, fertility, and developmental timing (True 2003; Wittkopp and Beldade 2009; Heath *et al*. 2013; Lindstedt *et al*. 2016). Temporal and spatial variation in these diverse selection pressures likely contribute to the abundant color variation in the genus *Neodiprion*.

As a first step to understanding the proximate mechanisms that generate color variation in pine sawflies, we conducted a quantitative trait locus (QTL) mapping study of larval body color and larval spotting pattern in the redheaded pine sawfly, *Neodiprion lecontei*. This species is widespread across eastern North America, where it feeds on multiple pine species. Throughout most of this range, larvae have several rows of dark black spots overlaid on a bright, yellow body (*e.g*., center image in Figure 1). However, previous field surveys and demographic analyses indicate that there has been a loss of yellow pigmentation in some populations in the Mid-Atlantic/northeastern United States and a pronounced reduction in spotting in populations living in the Great-Lakes region of the U.S. and Canada (Bagley *et al*. 2017; Figure 2). To describe genetic architectures and identify candidate genes underlying these two reduced-pigmentation phenotypes, we crossed white-bodied, dark-spotted individuals from a Virginia population to yellow-bodied, light-spotted individuals from a Michigan population (Figure 2). After mapping larval body-color and spotting-pattern traits in recombinant F_2_ haploid males, we used our linkage map to anchor the current *N. lecontei* genome assembly and identified candidate color genes within our QTL intervals. We conclude by comparing these initial findings on color genetics in pine sawflies with published studies from other insect taxa.

**Figure 2.**
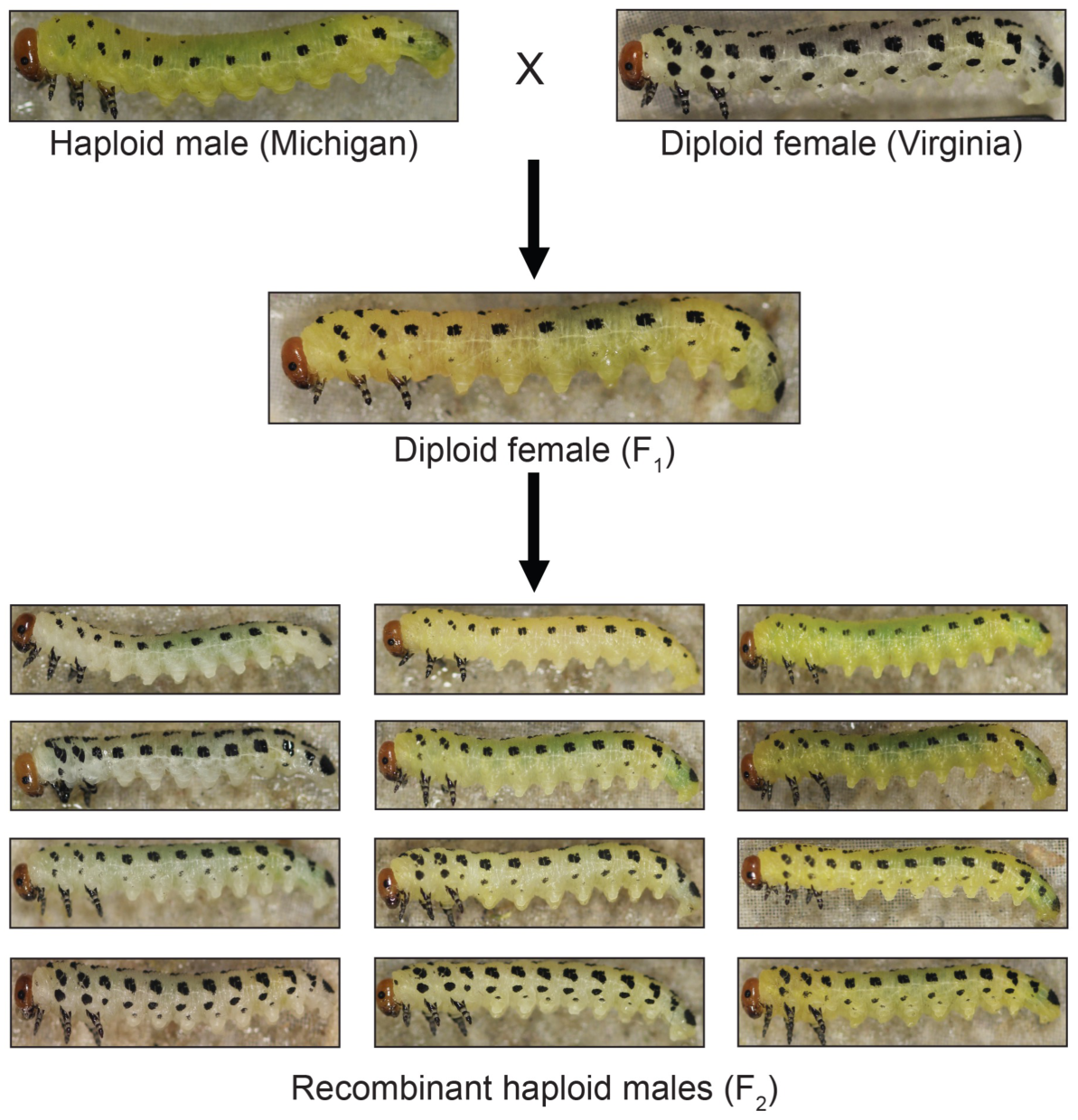
Intraspecific variation in *Neodiprion lecontei* larval color and cross design. We crossed white, dark-spotted diploid females from Virginia to yellow, light-spotted haploid males from Michigan. This produced haploid males with the VA genotype and phenotype (not shown) and diploid females (F_1_) with intermediate spotting and color. Virgin F_1_ females produced recombinant haploid males (F_2_) with a wide range of body color and spotting pattern (a representative sample is shown).

## MATERIALS AND METHODS

### Cross Design

To investigate the genetic basis of sawfly color traits, we crossed *Neodiprion lecontei* females from a white-bodied, dark-spotted population (collected from Valley View, VA; 37°54’47”N, 79°53’46”W) to *N. lecontei* males from a yellow-bodied, light-spotted population (collected from Bitely, MI; 43°47’46”N, 85°44’24”W). Both populations were collected from the field in 2012 and reared on *Pinus banksiana* (jack pine) for at least two generations in the lab via standard rearing protocols (described in more detail in Harper *et al*. 2016; Bendall *et al*. 2017). Our mapping families were derived from four grandparental pairs, which produced 10 F_1_ females. Like most hymenopterans, *N. lecontei* adults reproduce via arrhenotokous haplodiploidy, in which unfertilized eggs develop into haploid males and fertilized eggs develop into diploid females (Heimpel and de Boer 2008; Harper *et al*. 2016). Therefore, to produce an F_2_ haploid generation, we allowed virgin F_1_ females to lay eggs and reared their haploid male progeny on *P. banksiana* foliage until they reached a suitable size for phenotyping.

### Color phenotyping

*N. lecontei* larvae pass through five (males) or six (females) feeding instars and a single non-feeding instar, which are distinguishable on the basis of color pattern and size (Benjamin 1955; Coppel and Benjamin 1965; Wilson *et al*. 1992). For phenotyping, we chose only mature feeding larvae, which have an orange-red head capsule with a black ring around each eye and up to four pairs of rows of black spots (Wilson *et al*. 1992). Immediately after phenotyping, we preserved each larva in 100% ethanol for molecular work. In total, we generated color-phenotype data for 30 individuals from the VA population (mixed sex), 30 individuals from the MI population (mixed sex), 47 F_1_ females, and 429 F_2_ males (progeny of 10 virgin F_1_ females).

### Larval body color

We quantified larval body color in two ways: spectrometry and digital photography. First, we immobilized larvae with CO_2_ and recorded 15 reflectance spectra—five measurements from each of three body regions: the dorsum, lateral side, and ventrum—using a USB2000 spectrophotometer with a PX-2 Pulsed Xenon light source and SpectraSuite software (Ocean Optics, Largo, FL). We then used the program CLR: Colour Analysis Programs v1.05 (Montgomerie 2008) to trim the raw reflectance data to 300-700nm, compute 1-nm bins, and to compute the following summary statistics (described in more detail in the CLR documentation): B1, B2, B3, S1R, S1G, S1B, S1U, S1Y, S1V, S3, S5a, S5b, S5c, S6, S7, S8, S9, H1, H3, H4a, H4b, and H4c. This procedure produced 15 estimates for each summary statistic for each larva, which we then averaged to obtain a single value per statistic per larva. To reduce the 22 color summary statistics to a smaller number of independent variables, we performed a principal component analysis (PCA). To ensure that each variable was normally distributed, we performed a normal-quantile transformation prior to performing the PCA. Based on examination of the resulting scree plot, we retained the first two principal components (PC1 and PC2), which together accounted for 73.7% of the variance in the spectral data. Based on factor loadings, PC1 (PVE: 48.2%) corresponds to saturation (S1G, S1Y, S8) and brightness (B1, B2), while PC2 (PVE: 25.5%) corresponds to saturation (S5A, S5B, S5C, S6) and hue (H4c) (Table S1). Unless otherwise noted, the PCA and all other statistical analyses were performed in R (Version 3.1.3 or 3.2.3).

Because it provides objective and information-rich data, spectrometry is generally considered to be the gold standard for color quantification (Endler 1990; Andersson and Prager 2006). However, despite efforts to keep background lighting and reflectance probe positioning as consistent as possible across samples, measurement of small patches of yellow on rounded larval bodies proved challenging. Given the potential for variation in reflectance probe position to introduce substantial noise into our data (Montgomerie 2006), we used digital photography as a complementary method for quantifying larval body color (Stevens *et al*. 2007). For this analysis, we photographed CO2-immobilized larvae (dorsal and lateral surfaces) with a Canon EOS Rebel t3i camera equipped with an Achromat S 1.0X FWD 63mm lens and mounted on a copy stand. All photographs were taken in a dark, windowless room, and larvae were illuminated by two SLS CL-150 copy lamp lights, each with a 23-Watt (100W equivalent) soft white compact fluorescent light bulb. We then used Adobe Photoshop CC 2014 or 2015 (Adobe Systems Incorporated, San Jose, CA) to ascertain the amount of yellow present, following O’Quin *et al*. (2013). First, we converted each digital image (lateral surface) to CMYK color mode. Next, we selected the eye-dropper tool (set to a size of 5x5 pixels) as the color sampler tool, which we used to sample three different body locations: the body just behind the head and parallel to the eye, the first proleg, and the anal proleg. For each of the three regions, this procedure yielded an estimate of the proportion of the selected area that was yellow. We then averaged the three measurements to produce a single final measurement of yellow pigmentation (hereafter referred to as “yellow”).

### Larval spotting pattern

On each side, *N. lecontei* larvae have up to four rows of up to 11 spots extending from the mesothorax to the 9^th^ abdominal segment (Wilson *et al*. 1992). We quantified the extent of this black spotting in two ways: the number of spots present (spot number) and the proportion of body area covered by spots (spot area). To quantify spot number, we simply counted the number of spots in digital images of the dorsal and lateral surfaces (one side only). To quantify larval spotting area, we used Adobe Photoshop’s quick-selection tool to measure the area of the larval body (minus the head capsule) and the area of each row of lateral black spots. To control for differences in larval size, we divided the summed area of all lateral black spots by the area of the larval body. We also used the larval images and Adobe Photoshop to calculate the area of the head capsule, which we used as a covariate in some analyses to control for larval size (see below). We used a custom Perl script to process Photoshop measurement output files in bulk (written by John Terbot II; available in File S3).

### Statistical analysis of phenotypic data

In total, we produced three measures of body color (PC1, PC2, and yellow) and two measures of spotting pattern (spot number and spot area). To determine whether mean phenotypic values for the five color traits differed between the two populations and among the three generations of our cross, we performed Welch’s two-tailed *t*-tests, followed by Bonferroni correction for multiple comparisons. To determine the extent to which these five traits co-varied in the F_2_ males, we calculated Pearson’s correlation coefficients (*r*) and associated *P*-values, followed by Bonferroni correction for multiple comparisons. To determine which covariates to include in our QTL models, we performed ANOVAs to evaluate the relationship between each phenotype in the F_2_ males (429 total) and their F_1_ mothers (10 total) and head capsule sizes (a proxy for larval size/developmental stage). These and all other statistical analyses were performed in R version 3.3.2 (R Core Team 2013)

### Genotyping

We extracted DNA from ethanol-preserved larvae using a modified CTAB method (Chen *et al*. 2010) and prepared barcoded and indexed double-digest RAD (ddRAD) libraries using methods described elsewhere (Peterson *et al*. 2012; Bagley *et al*. 2017). We prepared a total of 10 indexed libraries: one consisting of the eight grandparents and 10 F_1_ females (18 adults total), and the remaining nine consisting of F_2_ haploid male larvae (~48 barcoded males per library). After verifying library quality using a Bioanalyzer 2100 (Agilent, Santa Clara, CA), we sent all 10 libraries to the University of Illinois Urbana-Champaign Roy J. Carver Biotechnology Center (Urbana, IL), where the libraries were pooled and sequenced using 100-bp single-end reads on two Illumina HiSeq2500 lanes. In total, we generated 400,621,900 reads.

We de-multiplexed and quality-filtered raw reads using the protocol described in Bagley *et al*. 2017. We then used Samtools v0.1.19 (Li *et al*. 2009) to map our reads to our *N. lecontei* reference genome (Vertacnik *et al*. 2016) and STACKS v1.37 (Catchen *et al*. 2013) to extract loci from our reference alignment and to call SNPs. We called SNPs in two different ways. First, for QTL mapping analyses, our goal was to recover fixed differences between the grandparental lines. To do so, we first called SNPs in our eight grandparents and 10 F_1_ mothers. For these 18 individuals, we required that SNPs had a minimum of 7x coverage and no more than 12% missing data. We then compiled a list of SNPs that represented fixed differences and, as an additional quality check, confirmed that all F_1_ females were heterozygous at these loci. We then used STACKS to call SNPs in the F_2_ haploid males, requiring that each SNP had a minimum of 5x coverage, no more than 10% missing data, and was present in the curated list produced from the grandparents. Filtering in STACKs produced 559 SNPs genotyped in 429 F_2_ males.

Second, to maximize the number of SNPs available for genome scaffolding, we performed an additional STACKS run using only the F_2_ haploid males, requiring that each SNP had a minimum of 4x coverage. By removing the requirement that SNPs were called in all grandparents, we could recover many more SNPs. We then filtered the data in VCFtools v0.1.14 (Danecek *et al*. 2011) to remove individuals with a mean depth of coverage less than one, retaining 408 F_2_ males. After removing low-coverage individuals, we used VCFtools to remove sites with a minor allele frequency (MAF) less than 0.05, sites with >5 heterozygotes (in a sample of haploid males, high heterozygosity is a clear indication of pervasive genotyping error), and sites with more than 50% missing data.

### Linkage map construction and genome scaffolding

To construct a linkage map for interval mapping, we started with 559 SNPs scored in 429 F_2_ males. After an additional round of filtering in R/qtl (Broman and Sen 2009), we removed 11 haploid males that had >50% missing data, for a total of 418 F_2_ males. Additionally, after removing SNPs that were genotyped in <70% of individuals, had identical genotypes to other SNPs, and had distorted segregation ratios (at α<0.05, after Bonferroni correction for multiple testing), we retained 503 SNPs. To assign these markers to linkage groups, we then used the “formLinkageGroups” function, requiring a minimum logarithm of odds (LOD) score of 6.0 and a maximum recombination frequency of 0.35. To order markers on linkage groups, we used the “orderMarkers” function, with the Kosambi mapping function to allow for crossovers. Following this initial ordering, we performed rippling on each linkage group to check whether switching marker order could improve LOD scores.

Our initial map included 503 SNPs spread across 358 scaffolds (out of 4523 scaffolds; Vertacnik *et al*. 2016). To increase the number of scaffolds and bases that we could place on our linkage groups, we performed additional linkage mapping analyses with a larger SNP dataset that was called as described above. We then constructed a linkage map for each of our four grandparental families (N = 54, 73, 120, and 161). For each grandparental family, we first performed additional data filtering in R/qtl to remove duplicate SNPs and SNPs with distorted segregation ratios, retaining between 2,049 and 3,155 SNPs per family. We then used the “formLinkageGroups” command with variable LOD thresholds (range: 5-15) and a maximum recombination frequency of 0.35. Because SNPs were not coded according to grandparent of origin, many alleles were “switched”. We therefore performed an iterative process of linkage group formation, visualization of pairwise recombination fractions and LOD scores (“plotRF” command), and allele switching (“switchAlleles” command) until we obtained seven linkage groups (the number of *N. lecontei* chromosomes; Smith 1942; Maxwell 1958; Sohi and Ennis 1981) and a recombination/LOD plot indicative of linkage within, but not between, linkage groups. Because allele ordering and examination of alternative SNP orders for these larger panels of SNPs was prohibitively slow in R/qtl, we used the more efficient MSTmap algorithm, implemented in R/ASMap v0.4-7 (Taylor and Butler 2017), to order our markers along their assigned linkage groups.

Finally, to order and orient our genome scaffolds along linkage groups (chromosomes), we used ALLMAPS (Tang *et al*. 2015) to combine information from our five maps (initial map with all individuals, but limited markers; plus four additional maps, each with more markers, but fewer individuals). Because maps constructed from larger families are likely to be more accurate than those constructed from small families, we weighted the maps according to their sample sizes.

### Interval mapping analysis

After linkage map construction, we used R/qtl to map QTL for our five color traits. Based on analyses of phenotypic covariates (Table S2), we included F_1_ mothers and head-capsule size as covariates in our analyses of PC1, spot number, and spot area; F_1_ mothers as covariates in our analysis of PC2; and no covariates in in our analysis of yellow. For each trait, we performed interval mapping using multiple imputation mapping. We first used the “sim.geno” function with a step size of 0 (*i.e*., genotypes only drawn at marker locations) and 64 replicates. We then used the “stepwiseqtl” command to detect QTL and select the multiple QTL model that optimized the penalized LOD score (Broman and Sen 2009; Manichaikul *et al*. 2009). To obtain penalties for the penalized LOD scores, we used the “scantwo” function to perform 1,000 permutations under a two-dimensional, two-QTL model that allows for interactions between QTL and the “calc.penalties” function to calculate penalties from these permutation results, using a significance threshold of α = 0.05. Finally, for each QTL retained in the final model, we calculated a 1.5-LOD support interval.

### Candidate gene analysis

As a first step to moving from QTL intervals to causal loci, we compiled a list of candidate color genes and determined their location in the *N. lecontei* genome. For larval spotting, we included genes in the melanin synthesis pathway and genes that have been implicated in pigmentation patterning (Wittkopp *et al*. 2003; Protas and Patel 2008; Wittkopp and Beldade 2009; Sugumaran and Barek 2016). For larval body color, we included genes implicated in the transport, deposition, and processing of carotenoid pigments derived from the diet (Palm *et al*. 2012; Yokoyama *et al*. 2013; Tsuchida and Sakudoh 2015; Toews *et al*. 2017). Although several pigments can produce yellow coloration in insects (*e.g*., melanins, pterins, ommochromes, and carotenoids), we focused on carotenoids because a heated pyridine test (McGraw *et al*. 2005) was consistent with carotenoid-based coloration in *N. lecontei* larvae (Figure S1).

Once we had compiled a list of candidate genes, we searched for these genes by name in the *N. lecontei* v1.0 genome assembly and NCBI annotation release 100 (Vertacnik *et al*. 2016). To find missing genes and as an additional quality measure, we obtained FASTA files corresponding to each candidate protein and/or gene from NCBI (using *Apis, Drosophila melanogaster,* or *Bombyx mori* sequences, depending on availability). We then used the i5k Workspace@NAL (Poelchau *et al*. 2014) BLAST (Altschul *et al*. 1990) web application to conduct tblastn (for protein sequences) or tblastx (for gene sequences) searches against the *N. lecontei* v1.0 genome assembly, using default search settings. After identifying the top hit for each candidate gene/protein, we then used the WebApollo (Lee *et al*. 2013) JBrowse (Skinner *et al*. 2009) *N. lecontei* genome browser to identify the corresponding predicted protein coding genes (from NCBI annotation release 100) in the *N. lecontei* genome.

We took additional steps to identify genes in the *yellow* gene family, all of which contain a major royal jelly protein (MRJP) domain (Ferguson *et al*. 2011). First, we used the search string “major royal jelly protein *Neodiprion*” to search the NCBI database for all predicted *yellow*-like and *yellow-MRJP*-like *N. lecontei* genes. We then downloaded FASTA files for the putative *yellow* gene sequences (26 total). Next, we used the Hymenoptera Genome Database (Elsik *et al*. 2016) to conduct a blastx search of our *N. lecontei* gene sequence queries against the *Apis mellifera* v4.5 genome NCBI RefSeq annotation release 103. Finally, we recorded the top *A. mellifera* hit for each putative *N. lecontei yellow* gene.

### Data availability

Short-read DNA sequences will be made available via the NCBI SRA (Bioproject PRJNA#######, Biosample numbers SAMN########-SAMN########). The linkage-group anchored assembly will be submitted to NCBI and i5k to update the existing *N. lecontei* genome assembly and annotations (Vertacnik *et al*. 2016). File S3 contains the custom script used to process the raw spot-area data. File S4 contains phenotypic data from all generations. File S5 contains the input file for R/qtl. Files S6 and S7 contain the genotype data and linkage maps used in the scaffolding analysis, respectively.

## RESULTS AND DISCUSSION

### Variation in larval color traits

Lab-reared larvae derived from the two founding populations differed significantly from one another for all color phenotypes (Figures 2, 3; Table S3). Because all larvae were reared on the same host under the same laboratory conditions (*i.e*., minimal environmental variance), these results suggest that genetic variance contributes to variance in these larval color traits. Crosses between the VA and MI lines produced diploid F_1_ female larvae that were intermediate in both body color and spotting pattern (Figures 2, 3). With the exception of PC2, F_1_ larvae differed significantly from both grandparents for all color traits (Table S3), indicating a lack of complete dominance for these traits. In contrast, the yellow, MI-like hue/chroma (PC2) appears to be completely dominant to the white, VA-like hue/chroma. We can make additional inferences about trait dominance by comparing the phenotypes of diploid F_1_ female larvae (dominance effects present) to those of the haploid F_2_ male larvae (dominance effects absent). For all three body-color measures (yellow, PC1, and PC2), average F_1_ larval colors were more MI-like (yellow) than those of F_2_ larvae. In contrast, compared to the F_2_ larvae, spotting phenotypes (number and area) of F_1_ larvae are more similar to the VA population (dark-spotted). Taken together, these phenotypic data suggest that the more heavily pigmented body-color and spotting-pattern phenotypes (*i.e*., large values for yellow, spot number, and spot area; small values for PC1 and PC2) are partially dominant to the less pigmented phenotypes.

Larval color variation observed in the F_2_ males spanned—and even exceeded— the range of variation observed in the grandparental populations and F_1_ females (Figure 3). The observation that grandparental body-color and spotting-pattern phenotypes are recapitulated in the F_2_ males suggests that both types of traits are controlled by a relatively small number of loci. There are multiple, non-mutually exclusive explanations for the transgressive color phenotypes in our haploid F_2_ larvae, including: variation in the grandparental lines, reduced developmental stability in hybrids, epistasis, unmasking of recessive alleles in haploid males, and the complementary action of additive alleles from the two grandparental lines (Rieseberg *et al*. 1999).

**Figure 3.**
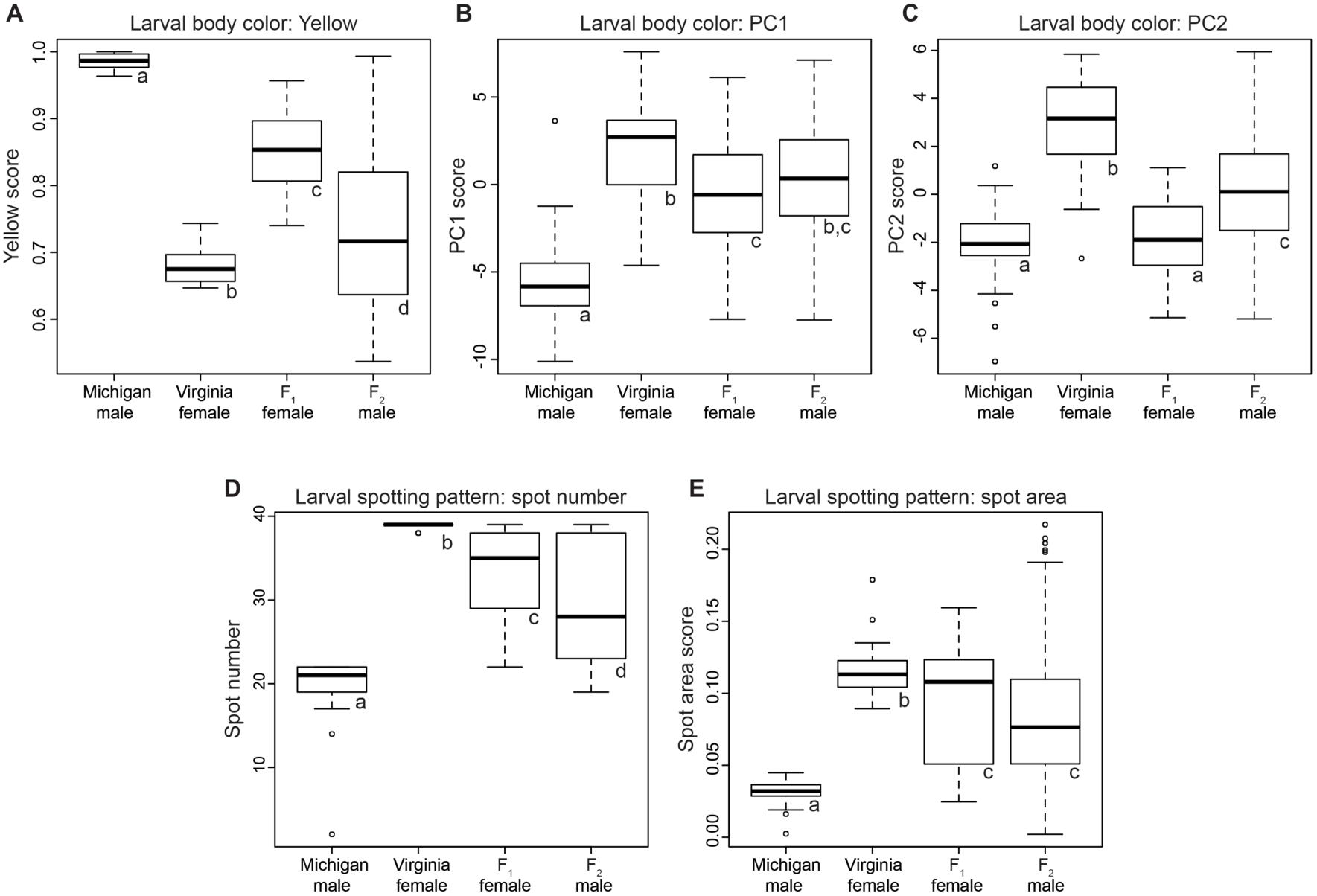
Larval color variation among *N. lecontei* populations and across generations. For larval body color (A-C), higher scores for yellow (A) and lower scores for PC1 (B) and PC2 (C) indicate higher levels of yellow pigment. For larval spotting pattern (D-E), higher spot numbers (D) and spot area scores (E) indicate more melanic spotting. For all traits, boxes represent interquartile ranges (median ± 2 s.d.), with outliers indicated as points, and letters denote significant differences after correction for multiple comparisons (see Table S3 for full statistical results).

We also observed significant correlations between many of the color traits (Figure S2). Encouragingly, estimates of body color derived from digital images (yellow) correlated significantly with both estimates derived from spectrometry (PC1 and PC2), suggesting that these independent data sources describe the same underlying larval color phenotype (*i.e*., amount of yellow pigment). Similarly, we observed strong correlations between the number and area of larval spots, indicating that both measures reliably characterize the extent of melanic spotting on the larval body. We also found that larvae with yellower bodies (small values for PC2) tended to be less heavily spotted (Figure S2). This observation suggests that body color and spotting pattern do not evolve completely independently of one another. Nevertheless, the correlation between these traits was weak and we observed many different combinations of spotting and pigmentation in the recombinant F_2_ males (Figure 2).

### Linkage mapping and genome scaffolding

Our 503 SNP markers were spread across seven linkage groups (LGs), which matches the number of *N. lecontei* chromosomes (Smith 1941; Maxwell 1958; Sohi and Ennis 1981). The total map length was 1169 cM, with an average marker spacing of 2.4 cM and maximum marker spacing of 24.3 cM (Table S4; Figure 4). Together, these results indicate that this linkage map is of sufficient quality and coverage for interval mapping. Additionally, with an estimated genome size of 340 Mb (estimated via flow cytometry; C. Linnen, personal observation), these mapping results yield a recombination density estimate of 3.43 cM/Mb. This recombination rate is lower than that observed in social hymenopterans, which have among the highest rates of recombination in eukaryotes (Wilfert *et al*. 2007). Nevertheless, this rate is on par with that reported in other (non-eusocial) hymenopterans, which lends support to the hypothesis that elevated recombination rates in eusocial hymenopteran species is a derived trait and possibly an adaptation to a social lifestyle (Gadau *et al*. 2000; Schmid-Hempel 2000; Crozier and Fjerdingstad 2001).

**Figure 4.**
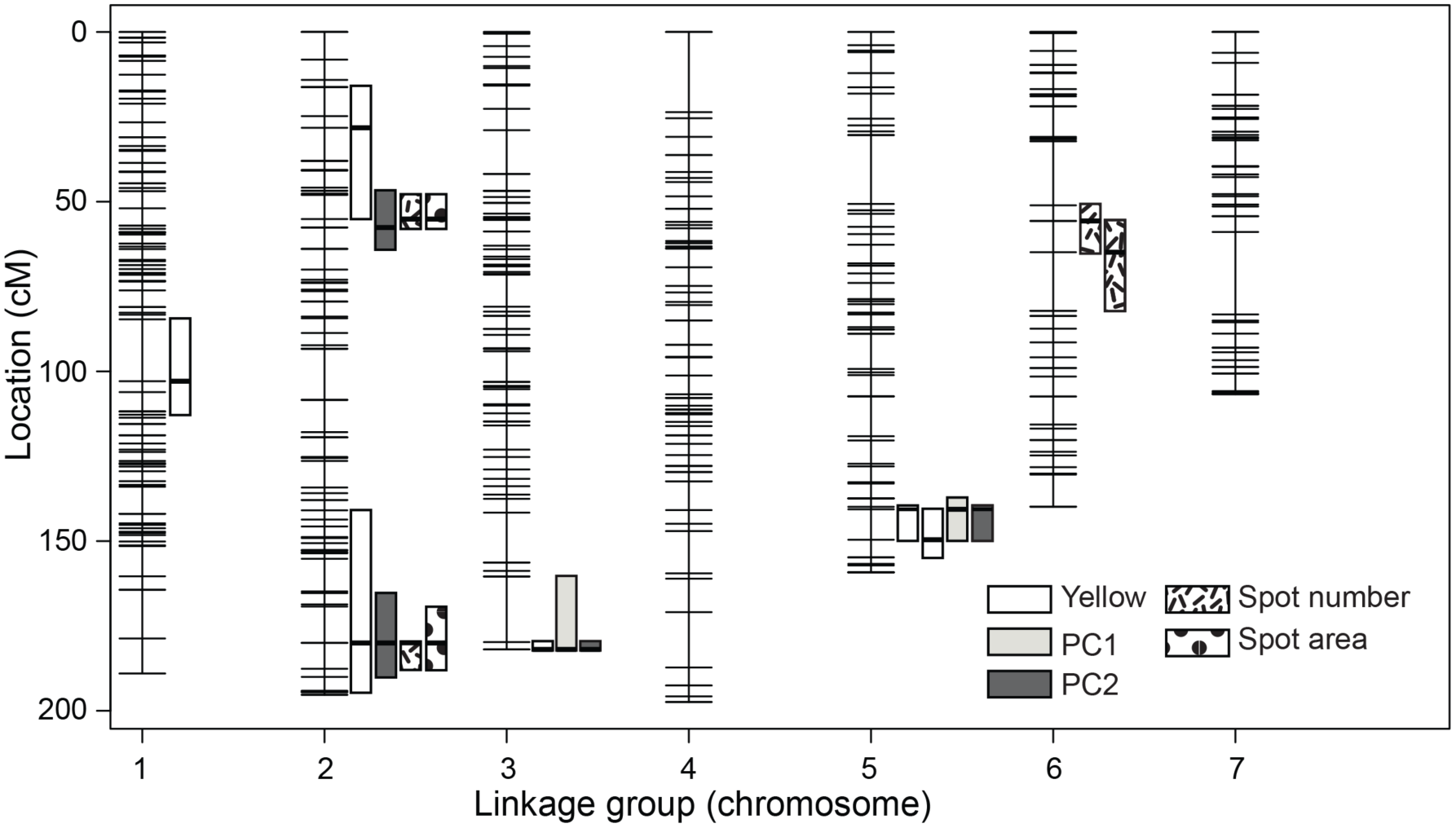
*N. lecontei* linkage map with larval-color QTL and 1.5-LOD support intervals.

Linkage maps estimated for the four grandparental families, each of which contained >2000 markers, ranged in length from 1072 cM to 3064 cM (Table S4). This variation in map length is likely attributable to both decreased mapping accuracy in smaller families and decreased genotyping accuracy in these less-stringently filtered SNP datasets. Nevertheless, our scaffolding analysis revealed that marker ordering was highly consistent across linkage maps (Figures S3-S9). Additionally, via including more SNPs, we were able to more than triple the number of mapped scaffolds (from 358 to 1005) and increase the percentage of mapped bases from 41.2% to 78.9% (Tables S5-S6).

Anchored genome scaffolds, coupled with existing *N. lecontei* gene annotations, are a valuable resource for identification of candidate genes within QTL.

### Genetic architectures of larval body color and larval spotting pattern

Combining the results from our three measures of larval body-color (yellow, PC1, and PC2), our interval mapping analyses detected a total of six distinct QTL regions and one QTL x QTL interaction (Table 1; Figures 4,5A). Of these, the largest-effect QTL were on LGs 3 and 5. We also recovered a significant interaction between these QTL for yellow (Figure 5B). Compared to analyses of larval body color, analyses of larval spotting pattern yielded fewer distinct QTL (four), but more QTL x QTL interactions (three) (Table 1; Figures 4, 5C, 5D). Both spotting measures (spot number and spot area) recovered two large-effect QTL on LG 2. These spotting QTL also overlapped with QTL with small-to-modest effects on body color (Table 1; Figure 4). Co-localization of spotting-pattern and body-color QTL suggests that the phenotypic correlations we observed between body color and spotting pattern in F_2_ males (Figure S2) are caused by underlying genetic correlations via either pleiotropy or physical linkage.

**Table 1.** QTL locations and effect sizes for larval body color (yellow, PC1, and PC2) and larval spotting pattern (spot number and spot area).

| Trait type | Trait | QTL name | LG<sup>4</sup> | Position (interval)<sup>5</sup> | Marker<sup>6</sup> | LOD | PVE | Effect size (SE)<sup>7</sup> | % diff.<sup>8</sup> |
| --- | --- | --- | --- | --- | --- | --- | --- | --- | --- |
| Body color | Yellow | Yellow-1 | 1 | 105.9 (84.4-112.5) | 5744 | 8.29 | 1.32 | -0.025 (0.0043) | 8.06 |
| Body color | Yellow | Yellow-2 | 2 | 28.2 (16.2-55.1) | 19162 | 2.92 | 0.45 | -0.014 (0.0043) | 4.63 |
| Body color | Yellow | Yellow-3 | 2 | 179.9 (140.8-194.0) | 15279 | 3.51 | 0.54 | -0.016 (0.0043) | 5.13 |
| Body color | Yellow | Yellow-4 | 3 | 181.9 (179.8-181.9) | 1882 | 71.23 | 16.27 | -0.084 (0.0043) | 27.19 |
| Body color | Yellow | Yellow-5 | 5 | 140.6 (139.9-149.6) | 3222 | 71.17 | 16.25 | -0.16 (0.0081) | 51.64 |
| Body color | Yellow | Yellow-6 | 5 | 149.6 (140.6-154.8) | 19661 | 4.37 | 0.68 | -0.035 (0.0082) | 11.43 |
| Body color | Yellow |  | Interaction | Yellow-3 x Yellow-5 |  | 12.45 | 2.03 | 0.061 (0.0086) | 19.58 |
| Body color | PC1 | PC1-1 | 3 | 181.9 (160.4-181.9) | 1882 | 4.31 | 3.56 | 1.15 (0.26) | 15.51 |
| Body color | PC1 | PC1-2 | 5 | 140.6 (137.4-149.6) | 3222 | 14.75 | 12.92 | 2.17 (0.26) | 29.30 |
| Body color | PC2 | PC2-1 | 2 | 55.1 (46.8-63.9) | 21 | 11.37 | 6.42 | 1.18 (0.16) | 23.46 |
| Body color | PC2 | PC2-2 | 2 | 179.8 (165.2-189.9) | 2143 | 7.24 | 3.99 | 0.91 (0.16) | 17.99 |
| Body color | PC2 | PC2-3 | 3 | 181.9 (179.8-181.9) | 1882 | 14.60 | 8.39 | 1.33 (0.16) | 26.39 |
| Body color | PC2 | PC2-4 | 5 | 140.6 (139.9-149.6) | 3222 | 33.13 | 21.13 | 2.09 (0.16) | 41.45 |
| Spotting pattern | Spot number | SpotNum-1 | 2 | 55.1 (48.0-57.6) | 9282 | 58.46 | 23.52 | 6.31 (0.36) | 33.06 |
| Spotting pattern | Spot number | SpotNum-2 | 2 | 179.9 (179.8-187.5) | 15279 | 78.19 | 35.45 | 7.81 (0.35) | 40.96 |
| Spotting pattern | Spot number | SpotNum-3 | 6 | 55.7 (51.1-64.9) | 9040 | 4.84 | 1.43 | -0.58 (0.59) | 3.05 |
| Spotting pattern | Spot number | SpotNum-4 | 6 | 64.9 (55.7-82.2) | 7485 | 6.75 | 2.02 | -1.00 (0.59) | 5.26 |
| Spotting pattern | Spot number |  | Interaction | SpotNum-1 x SpotNum-4 |  | 6.15 | 1.84 | 3.45 (0.71) | 18.09 |
| Spotting pattern | Spot number |  | Interaction | SpotNum-2 x SpotNum-3 |  | 4.64 | 1.38 | -3.13 (0.71) | 16.39 |
| Spotting pattern | Spot area | SpotArea-1 | 2 | 55.1 (48.0-57.6) | 9282 | 28.62 | 12.99 | 0.029 (0.0025) | 34.86 |
| Spotting pattern | Spot area | SpotArea-2 | 2 | 179.9 (169.1-187.5) | 15279 | 63.73 | 35.47 | 0.048 (0.0024) | 57.04 |
| Spotting pattern | Spot area |  | Interaction | Spot-1 x Spot-2 |  | 1.76 | 0.69 | 0.014 (0.0049) | 16.49 |

<sup>4</sup> Linkage group number
<sup>5</sup> Position in cM (1.5-LOD support intervals)
<sup>6</sup> Marker closest to QTL peak
<sup>7</sup> Effect sizes calculated as the difference in the phenotypic averages of among F<sub>2</sub> males carrying a VA allele and F<sub>2</sub> males carrying a MI allele ( ± standard error).
<sup>8</sup> Effect sizes calculated as a percentage of the difference between average trait values for the two grandparental lines (VA and MI).

**Figure 5.**
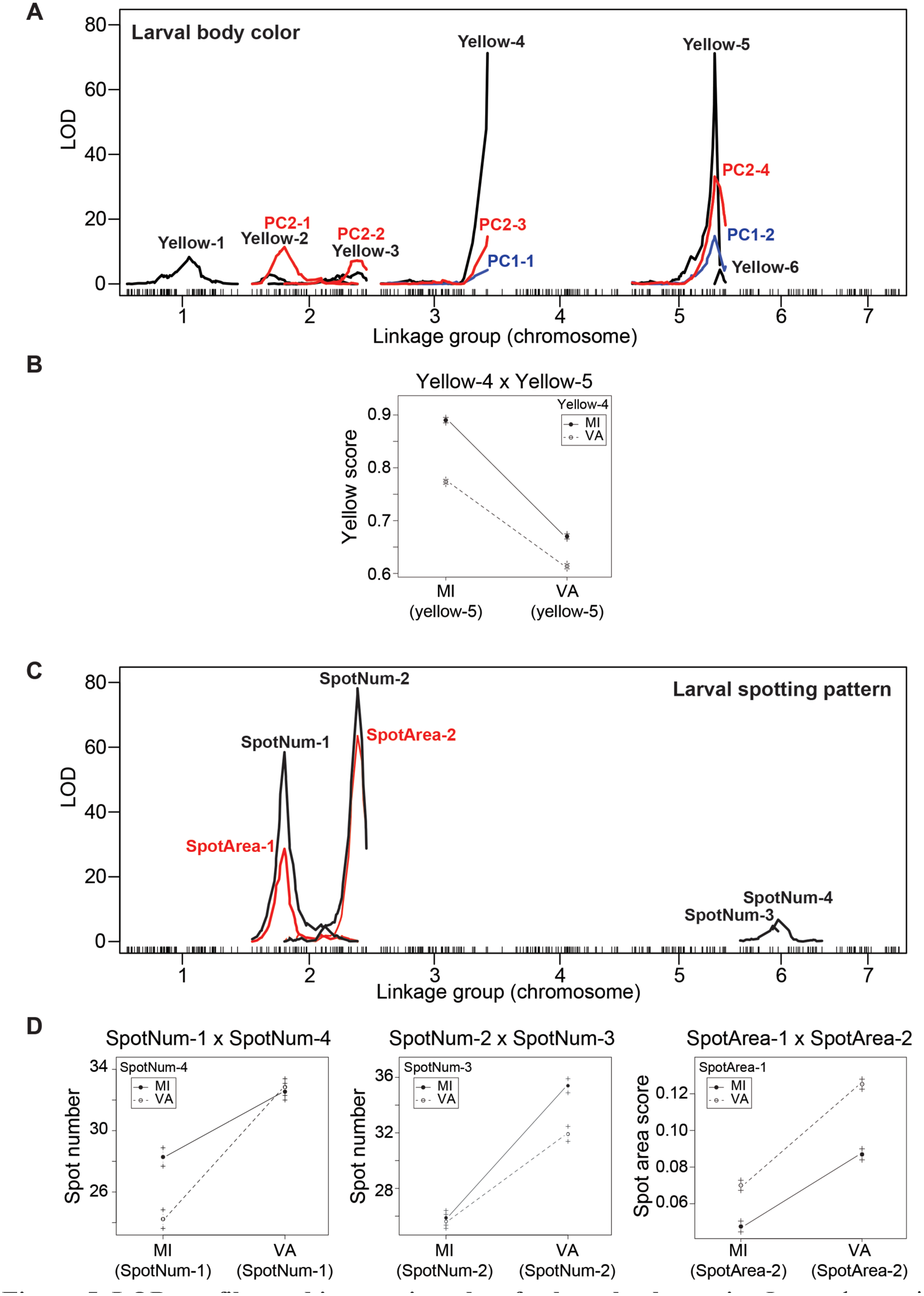
LOD profiles and interaction plots for larval color traits. Interval mapping analyses recovered QTL for larval body color (Yellow, PC1, PC2) on LGs 1, 2, 3, and 5 (A) and a single QTL x QTL interaction (B). QTL for larval spotting pattern (spot area and spot number) were on LGs 2 and 6 (C), with three QTL x QTL interactions (D). QTL names are as in Table 1.

These genetic mapping results provide us with our first glimpse into the genetic architecture of naturally occurring color variation in pine sawflies. For both larval body color and larval spotting pattern, we recovered a relatively small number of loci, some of which had a substantial effect on the color phenotype (Table 1). Additionally, our multiple-QTL models explained a considerable percentage of the observed variation in larval color: up to 86% for larval body color (yellow) and up to 73% for larval spotting pattern (spot number) (Table 2). Although limited statistical power precluded us from quantifying the number of QTL of very small effect, our results suggest that the proportion of variation explained by small, isolated QTL is small. It is possible, however, that our large-effect QTL comprise many linked QTL of individually small effect (*e.g*., Stam and Laurie 1996; McGregor *et al*. 2007; Bickel *et al*. 2011; Linnen *et al*. 2013). Distinguishing between oligogenic architectures (a small number of moderate to large-effect loci) and polygenic architectures (a large number of small-effect loci) will require fine-mapping and functionally validating genes and mutations for both color traits. Nevertheless, for both types of color traits, our results clearly rule out both a monogenic architecture and an architecture in which there are many unlinked, small-effect mutations. The latter architecture is also highly unlikely if local color adaptation in pine sawflies proceeded in the face of gene flow (Griswold 2006; Yeaman and Whitlock 2011), as previous demographic analyses seem to indicate is the case (Bagley *et al*. 2017).

**Table 2.** Summary of full multiple-QTL models for five larval-color traits.

| Trait type | Trait | Covariates | # QTL | # Interactions | LOD | PVE |
| --- | --- | --- | --- | --- | --- | --- |
| Body color | Yellow | None | 6 | 1 | 182.03 | 85.83 |
| Body color | PC1 | Size, Mother | 2 | 0 | 27.14 | 25.48 |
| Body color | PC2 | Mother | 4 | 0 | 65.84 | 50.92 |
| Spotting pattern | Spot number | Size, Mother | 4 | 2 | 122.18 | 73.23 |
| Spotting pattern | Spot area | Size, Mother | 2 | 1 | 94.77 | 63.93 |

To date, QTL-mapping and candidate-gene studies of color traits have yielded many examples of relatively simple genetic architectures in which one or a small number of mutations have very large phenotypic effects (Rockman 2012; Martin and Orgogozo 2013). However, the relevance of these studies to broader patterns in phenotypic evolution remains unclear due to biases that stem from a tendency to work on discrete phenotypes and specific candidate genes. These biases can be minimized by focusing on continuously varying color traits and employing an unbiased mapping approach as we have done here (see also O’Quin *et al*. 2013; Albertson *et al*. 2014; Signor *et al*. 2016; Yassin *et al*. 2016). Experimental biases aside, some have also argued that color itself— but not color pattern—is atypically genetically simple (i.e., more likely to be mono‐ or oligogenic than other traits) (Charlesworth *et al*. 1982; Rockman 2012). In our own data, we see no obvious differences between the architectures of body color and spotting pattern. If anything, the architecture of spotting appears slightly “simpler” (fewer QTL with larger individual effects) than that of body color (Table 1). That said, more spotting variation remains unexplained (Table 2), which could be attributable to a plethora of undetected small-effect loci, and thus a more complex underlying architecture. A more definitive test of the “color/pattern” hypothesis will therefore require fine-mapping our QTL, with the prediction that the spotting QTL will “fractionate” to a greater extent (i.e., break into more QTL of individually smaller effect) than the body-color QTL. A rigorous evaluation of the color/pattern hypothesis will also require data on the genetic architecture of color and pattern traits that have evolved independently in other taxa. To this end, ample variation in both body-color and color-pattern traits make *Neodiprion* sawflies an ideal system for determining whether there are predictable differences in the genetic architecture of different types of color traits.

### Candidate genes for larval color traits

In total, we identified 61 candidate genes with known or suspected roles in melanin-based or carotenoid-based pigmentation and for which we were able to identify putative homologs in the *N. lecontei* genome (Table S7). Of these, 26 appeared to belong to the *yellow* gene family. Notably, this number is equivalent to the number of *yellow* - like/*yellow-MRJP*-like genes found in the genome of the jewel wasp, *Nasonia vitripennis*, which boasts the highest reported number of *yellow*-like genes of any insect to date (Werren *et al*. 2010). Additionally, 13 of these genes (*yellow-e, yellow-e3*, four *yellow-g, yellow-h, yellow-x*, and five*MJRPs*) were found in tandem array along three adjacent scaffolds (548, 170, and 36; ~1 Mb total) on LG 2. This genomic organization is consistent with a conserved clustering of *yellow-h, ‐e3, ‐e, ‐g2*, and –*g* observed across *Apis, Tribolium, Bombyx, Drosophila,* and *Nasonia* (Drapeau *et al*. 2006; Werren *et al*. 2010; Ferguson *et al*. 2011). Like *Nasonia* and *Apis,* this cluster also contains *MJRPs;* like *Heliconius,* this cluster contains a *yellow-x* gene. Overall, we placed 57 of our 61 color genes on our LG-anchored assembly. With these data, we identified candidate genes for all but two QTL (Yellow-1 on LG 1 and SpotNum-3 on LG 6) (Table S7).

### Candidate genes for larval body color

Both of our largest-effect body-color QTL regions contained promising candidate genes with known or suspected roles in carotenoid-based pigmentation (Table S7). First, within the LG-3 QTL region (Yellow-4, PC1-1, PC2-3; Table 1, Figure 4), we found a predicted protein-coding region in scaffold 518 with a high degree of similarity to the *Bombyx mori* Cameo2 scavenger receptor protein (e-value: 1 x 10^-18^; bitscore: 92.8). *Cameo2* encodes a transmembrane protein belonging to the CD36 family that has been implicated in the selective transport of the carotenoid lutein from the hemolymph to specific tissues (Sakudoh *et al*. 2010; Tsuchida and Sakudoh 2015). In the silkworm *Bombyx mori*, *Cameo2* is responsible for the “C mutant” phenotype, which is characterized by a combination of yellow hemolymph and white cocoons that arises as a consequence of disrupted transport of lutein from the hemolymph to the middle silk gland (Sakudoh *et al*. 2010; Tsuchida and Sakudoh 2015).

Second, within the LG-5 QTL region (Yellow-5, Yellow-6, PC1-2, PC2-4; Table 1, Figure 4), we recovered a predicted protein coding gene in scaffold 164 with a high degree of similarity to the ApoLTP-1 and ApoLTP-2 protein subunits (encoded by the gene *apoLTP-II*/I) of the *Bombyx mori* lipid transfer particle (LTP) lipoprotein (e-value: 0; bitscore: 391). LTP is one of two major lipoproteins present in insect hemolymph and appears to be involved in the transport of hydrophobic lipids (including carotenoids) from the gut to the other major lipoprotein, lipophorin, which then transports lipids to target tissues (Tsuchida *et al*. 1998; Palm *et al*. 2012; Yokoyama *et al*. 2013). Based on these observations, we hypothesize that loss-of-function mutations in both *Cameo2* and *apoLTP-II*/I contribute to the loss of yellow pigmentation in white-bodied *N. lecontei* larvae.

### Candidate genes for larval spotting pattern

Our two largest-effect spotting-pattern QTL also yielded promising candidate genes—this time in the well-characterized melanin biosynthesis pathway (Table S7). In both LG-2 QTL regions, we found protein-coding genes that belong to the *yellow* gene family. The first LG-2 QTL region (SpotNum-1 and SpotArea-1; Table 1, Figure 4) contained 2 *yellow*-like genes that were most similar to *Apis yellow-x1* (e-value: 1.2 x 10^-160^; bitscore: 471.47 and e-value: 3.9 x 10^-151^; bitscore: 448.36). At present, there is little known about the function of *yellow-x* genes, which appear to be highly divergent from other *yellow* gene families (Ferguson *et al*. 2011). The second LG-2 QTL region (SpotNum-2 and SpotArea-2; Table 1, Figure 4) contained a cluster of 13 *yellow* genes, including *yellow-e*. In two different mutant strains of *B. mori,* mutations in *yellow-e* produced a truncated gene product that results in increased reddish-brown pigmentation in the head cuticle and anal plate compared to wildtype strains (Ito *et al*. 2010). In wildtype larvae, *yellow-e* is most highly expressed in the integument of the head and the tail (Ito *et al*. 2010). Based on these observations, one possible mechanism for the reduced spotting observed in the light-spotted MI population is an increase in *yellow-e* expression.

The second LG-2 spotting QTL region also contained a predicted protein that was highly similar to tyrosine hydroxylase (TH) (e-score: 6 x 10^-123^, bitscore: 406). TH catalyzes the hydroxylation of tyrosine to 3,4-dihydroxyphenylalanine (DOPA), a precursor to melanin-based pigments (Wright 1987). Work in the swallowtail butterfly *Papilio xuthus* and the armyworm *Pseudaletia separata* demonstrates that TH and another enzyme, dopa decarboxylase (DDC), are expressed in larval epithelial cells containing black pigment (Futahashi and Fujiwara 2005; Ninomiya and Hayakawa 2007). Furthermore, inhibition of either enzyme prevented the formation of melanin-based larval pigmentation patterns (Futahashi and Fujiwara 2005). Thus, a reduction in the regional expression of *TH* (aka *pale)* is another plausible mechanism underlying reduced spotting in the light-spotted MI population.

We also found candidate genes in some of our QTL of more modest effect. As noted above, the Yellow-2 and Yellow-3 QTL overlap with the two large-effect spotting QTL (Table 1, Figures 4,5). One possible explanation for this observation is that in addition to impacting spotting phenotypes, genes in the melanin biosynthesis pathway also impact overall levels of melanin throughout the integument and therefore overall body color. Finally, the SpotNum-4 QTL contained a gene encoding dopamine N-acetyl transferase *(Dat)*. Dopamine N-acetyl transferase, which catalyzes the reaction between dopamine and N-acetyl dopamine and ultimately leads to the production of a colorless pigment, is responsible for a difference in pupal case color between two *Drosophila* species (Ahmed-Braimah and Sweigart 2015).

### Comparison with color loci in other taxa

Identification of candidate genes within our QTL peaks provides an opportunity to compare our results with those from different insect taxa. For melanin-based traits, the bulk of existing genetic studies involve *Drosophila* fruit flies (Massey and Wittkopp 2016), with a handful of additional studies in wild and domesticated lepidopterans (*e.g*., Ito *et al*. 2010; Hof *et al*. 2016; Nadeau *et al*. 2016). A recent synthesis of *Drosophila* studies suggests that cis-regulatory changes in a handful of genes controlling melanin biosynthesis (*yellow*, *ebony*, *tan*, *Dat*) and patterning (*bab1*, *bab2*, *omb*, *Dll*, and *wg*) explain much of the intra‐ and interspecific variation in pigmentation (Massey and Wittkopp 2016). Although our data cannot speak to the contribution of cis-regulatory versus protein-coding changes, we do recover some of the same melanin biosynthesis genes (*yellow* and *Dat*). By contrast, all of the *Drosophila* melanin patterning genes fell outside of our spotting-pattern QTL intervals (Table S7). While we cannot rule these genes out completely, our results suggest that none of them plays a major role in this intraspecific comparison. Whether the same is true of patterning differences within and between other *Neodiprion* species remains to be seen.

Compared to melanin-based pigmentation, much less is known about the genetic underpinnings of variation in carotenoid-based pigmentation, which is widespread in nature (Heath *et al*. 2013; Toews *et al*. 2017). Nevertheless, studies in the domesticated silkworm (Sakudoh *et al*. 2010; Tsuchida and Sakudoh 2015), salmonid fish (Sundvold *et al*. 2011), and scallops (Liu *et al*. 2015) collectively suggest that scavenger receptor genes such as our candidate *Cameo2* may be a common and taxonomically widespread source carotenoid-based color variation (Toews *et al*. 2017). Beyond evaluating patterns of gene reuse, extensive intra‐ and interspecific variation in larval body color across the genus *Neodiprion* (Figure 1) has the potential to provide novel insights into the molecular mechanisms underlying carotenoid-based pigmentation.

### Summary and Conclusions

Our study, which focuses on naturally occurring color variation in an undersampled life stage (larva), taxon (Hymenoptera), and pigment type (carotenoids), represents a valuable addition to the invertebrate pigmentation literature. Our results also have several implications for both hymenopteran evolution and color evolution. First, we provide the first recombination-rate estimate for the Eusymphyta, the sister group to all remaining Hymenoptera (*e.g*., bees, wasps, and ants) (Peters *et al*. 2017). Our estimate suggests that the exceptionally high recombination rates observed in some social hymenopterans represents a derived state, supporting the hypothesis that increased recombination is an adaptation for increasing genetic diversity in a colony setting (Gadau *et al*. 2000; Schmid-Hempel 2000; Crozier and Fjerdingstad 2001; Wilfert *et al*. 2007). Second, for both larval body color and spotting pattern, we can conclude that intraspecific variation is due neither to a single Mendelian locus nor to a large number of unlinked, small-effect loci. It remains to be seen, however, whether color, color pattern, and other types of phenotypes differ predictably in their genetic architectures (Rockman 2012; Dittmar *et al*. 2016). Third, our QTL intervals contain several key players in melanin-based and carotenoid-based pigmentation in other taxa, including *yellow*, *Dat*, and *Cameo2* (Massey and Wittkopp 2016; Toews *et al*. 2017). Although functional testing is needed to establish causal links, we can rule out a major role for several core melanin-patterning genes in this intraspecific comparison. Finally, although much work remains—including fine-mapping QTL, functional analysis of candidate genes, and genetic dissection of diverse color traits in additional *Neodiprion* populations and species—our rapid progression from phenotypic variation to strong candidate genes demonstrates the tremendous potential of this system for addressing fundamental questions about the genetic basis of color variation in natural populations.

## ACKNOWLEDGEMENTS

For assistance with collecting and maintaining sawfly colonies, we thank past members of the Linnen laboratory, especially Robin Bagley, Adam Leonberger, Kim Vertacnik, Mary Collins, and Aubrey Mojesky. For assistance with phenotypic data collection, we thank Aubrey Mojesky, Mary Collins, and Ismaeel Siddiqqi. For advice on linkage mapping in R/qtl, we thank Karl Broman and Kelly O’Quin. For constructive comments on earlier versions of this manuscript, we thank Danielle Herrig, Emily Bendall, Kim Vertacnik, Katie Peichel, and two anonymous reviewers. Funding for this research was provided by the National Science Foundation (DEB-1257739; CRL), the United States Department of Agriculture National Institute of Food and Agriculture (2016-67014-2475; CRL), the Academy of Finland (project no. 257581; CL), and the University of Kentucky (Ribble summer undergraduate research fellowship, honors program independent research grant, and summer research and creativity fellowship to TS).

